## Supplementary material for "DNA binding redistributes activation domain ensemble and accessibility in pioneer factor Sox2"

### Supplementary information

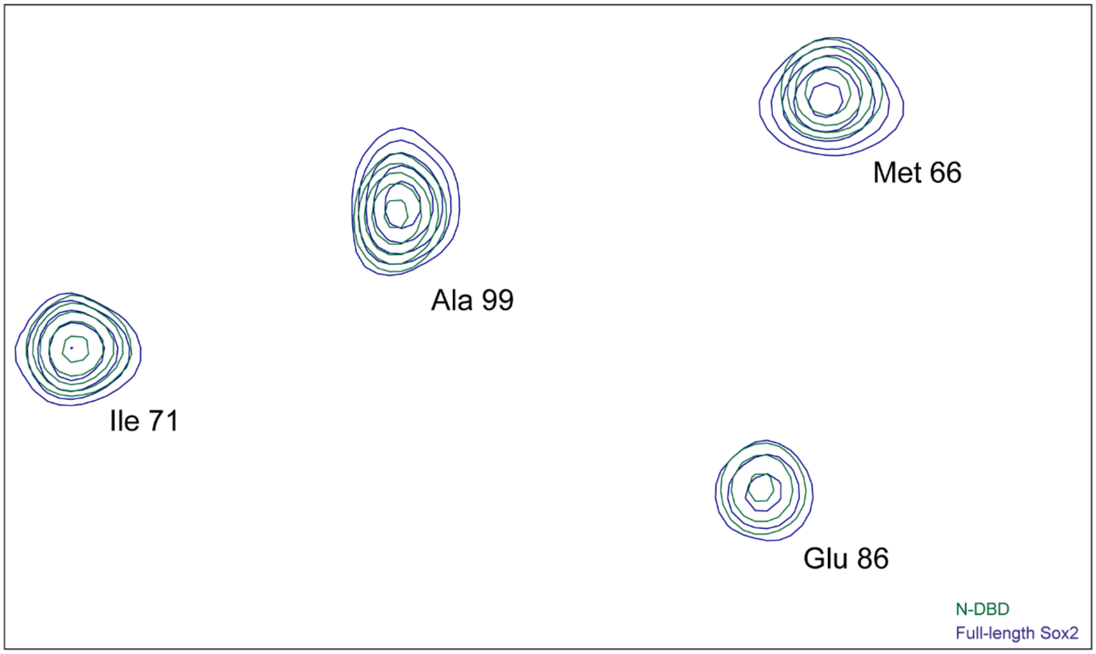


**Fig. S1. Peak positions and intensities of Sox2 DBD.** $^{1}H^{15}N$ BTROSY[1] of full-length Sox2 (blue) overlapped with $^{1}H^{15}N$ BTROSY of the N-DBD, showing that the peaks originating from the DBD overlap well with peaks from an isolated DBD. This indicates that the DBD structure is generally unperturbed by the presence of the C-IDR. Both spectra were run with 640 scans.

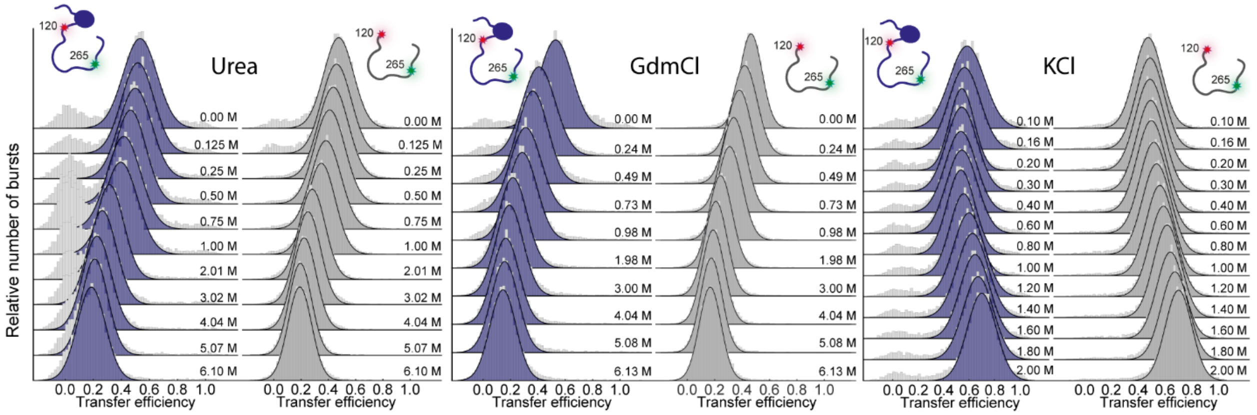


**Fig. S2. Dimensions of Sox2 C-IDR change in denaturants and salt.** Transfer efficiency histograms of full-length Sox2 (blue) or isolated C-IDR (grey) fluorescently labelled in positions 120-265, in different concentrations of urea (left), GdmCl (middle), and KCl (right).

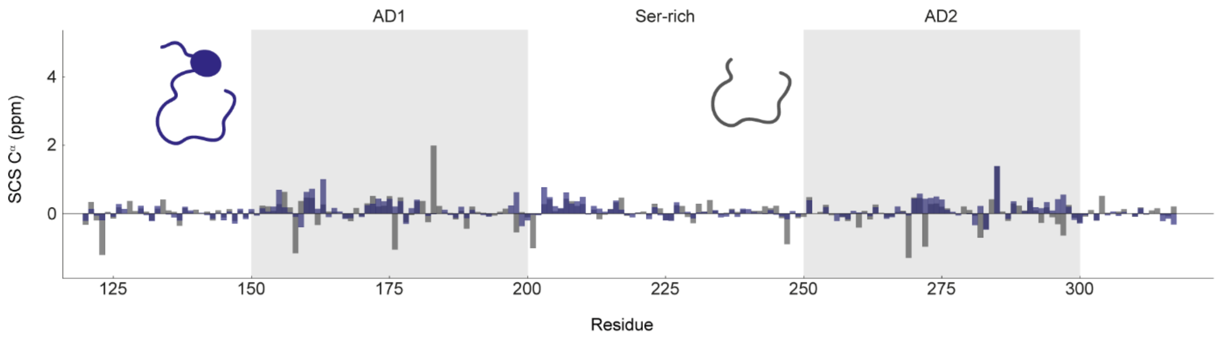


**Fig. S3. Cα SCS plot for residues 120-317 for full-length Sox2 and the isolated C-IDR.** Secondary structure content is very similar in the two constructs and indicates general lack of structure. The main domains are indicated.

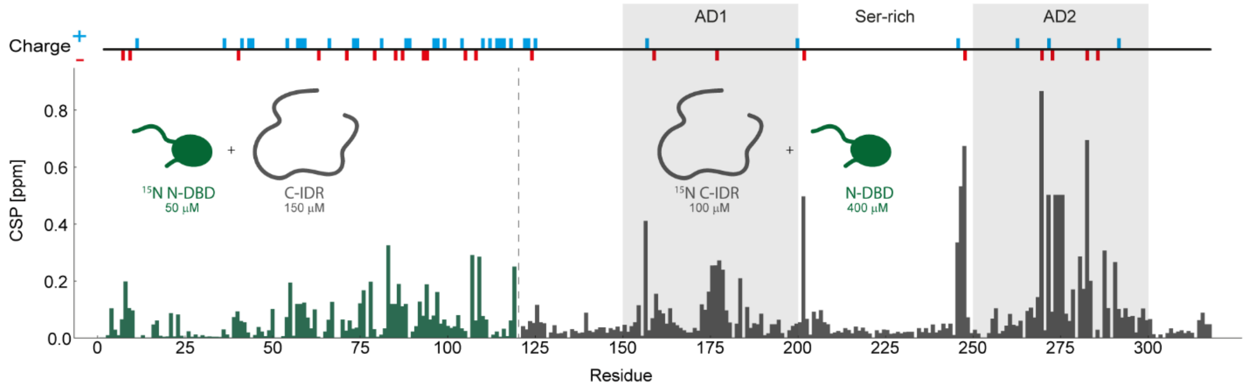


**Fig. S4. CSP plot of $^{15}$N-labelled individual domains mixed with their unlabelled counterpart domain.** The left side of the plot contains the combined $^{1}$H,$^{15}$N CSPs (see Methods) for the isolated $^{15}$N-labelled N-DBD (50 µM) with unlabelled C-IDR (150 µM). The right side of the plot contains the CSPs for the isolated $^{15}$N-labelled C-IDR (100 µM) with unlabelled N-DBD (400 µM).

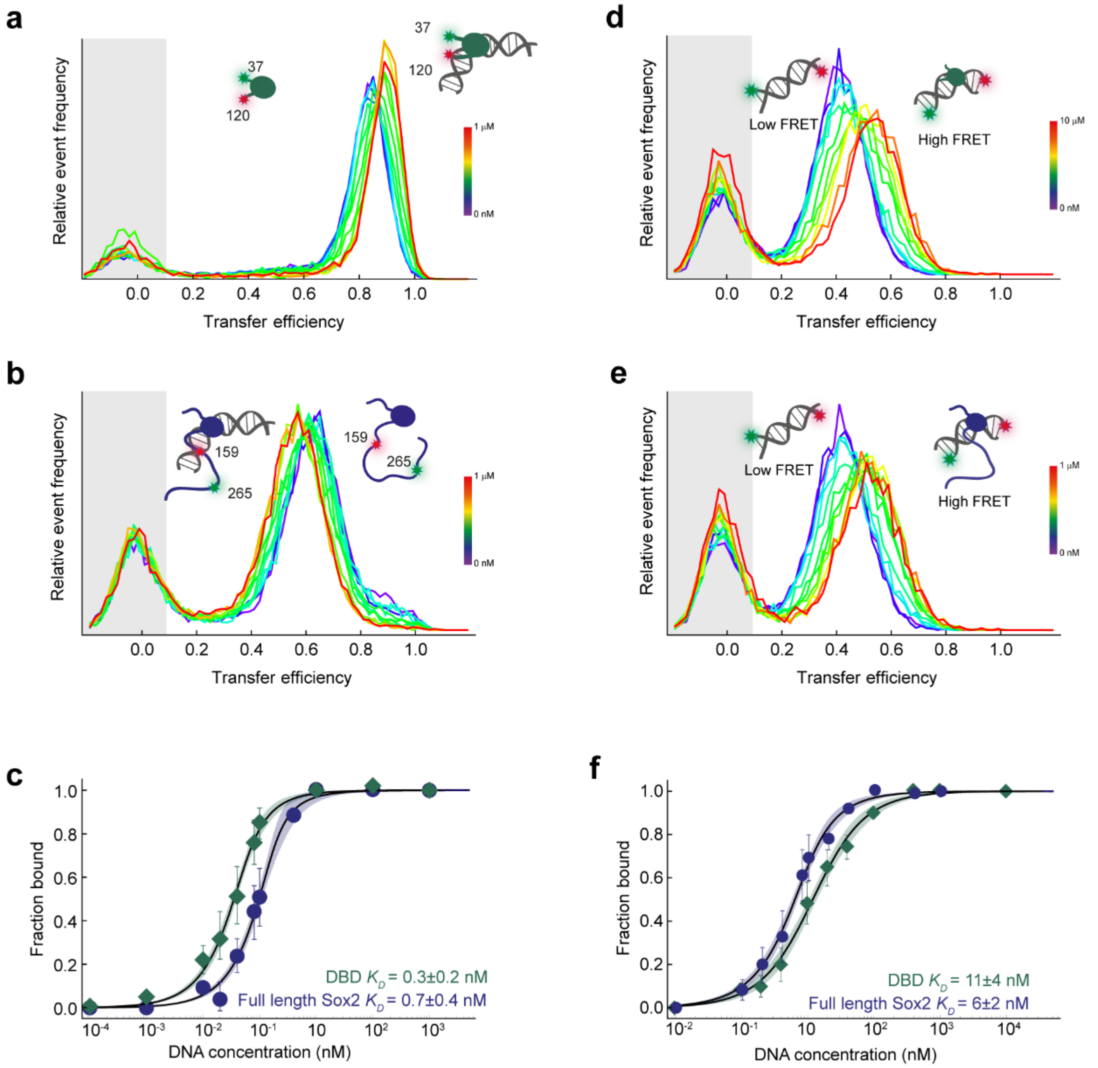


**Fig. S5. Binding affinity of Sox2 to specific and non-specific DNA. a-b)** Single-molecule transfer efficiency histograms of the a) isolated Sox2 DBD fluorescently labelled in positions 37 and 120 or b) full-length Sox2 fluorescently labelled in positions 159-265, with varying concentrations of unlabelled 30 bp specific DNA. **c)** The corresponding binding isotherms for panels a) and b). **d-e)** Single-molecule transfer efficiency histograms of 15 bp non-specific DNA fluorescently labelled at the 5´ and 3´-ends (see Supplementary Table 3) with varying concentrations of d) unlabelled isolated DBD or e) unlabelled full-length Sox2. **f)** The corresponding binding isotherms for panels d) and e). The dissociation constant for specific DNA using labelled proteins is nearly identical to the one determined with labelled DNA, excluding adverse effects from the fluorophores. Grey boxes in transfer efficiency histograms indicate donor-only populations, shaded areas in binding isotherms represent 95% confidence intervals of the fits.

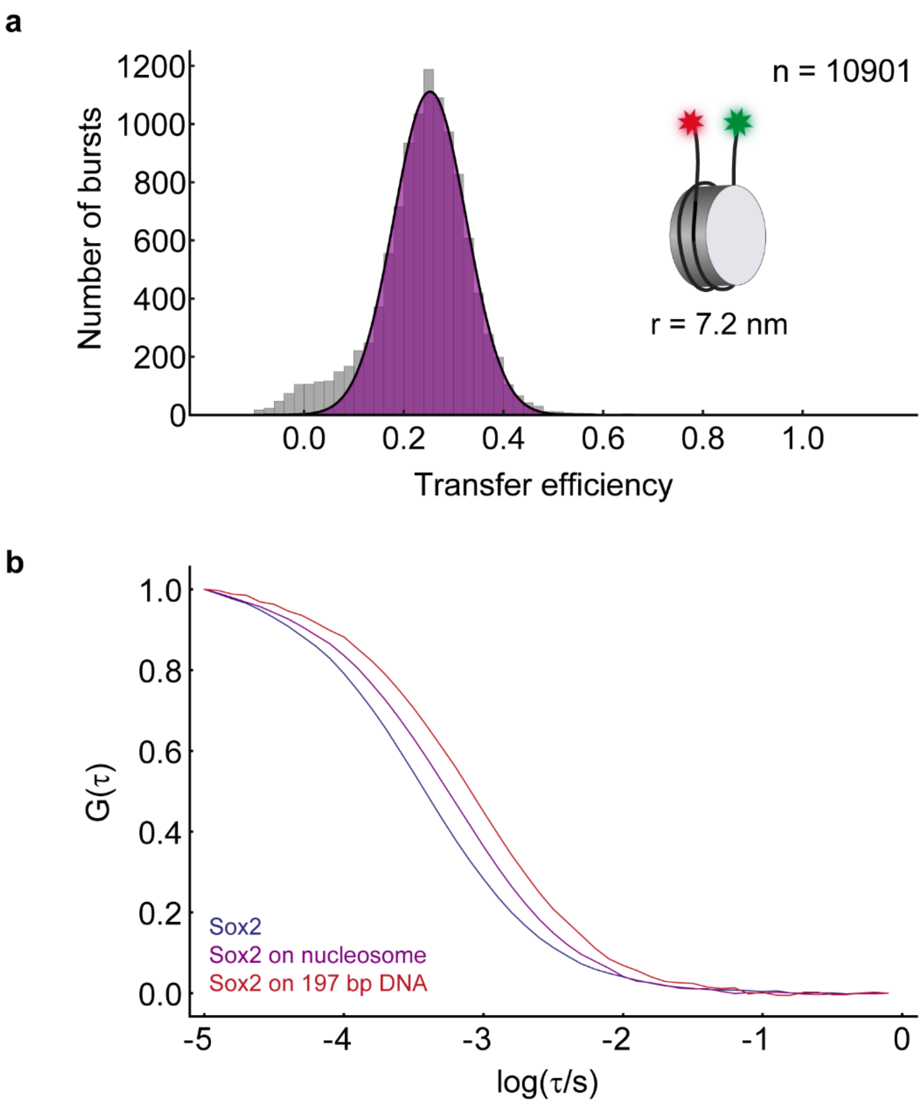


**Fig. S6. Single-molecule spectroscopy analysis of nucleosomes**. **a)** Single-molecule transfer efficiency histogram of 197 bp Widom 601 nucleosome fluorescently labelled at the DNA linker ends. Even at 100 pM concentrations, the nucleosome is stably wrapped as evident from the significant FRET between the fluorophores on each linker, in agreement with previous results[2]. The distance between the dyes is indicated, as calculated from the Förster equation. **b)** Donor-acceptor cross-correlation of fluorescently labelled Sox2, labelled in positions 120 and 315. Free Sox2 (blue) has a relatively short diffusion time through the confocal volume. In the presence of 90 nM of the same nucleosome as in panel a (unlabelled) the diffusion time is considerably increased (purple). In the presence of 197 bp 601 Widom DNA (red, without histone octamer), the diffusion time is increased even more, due to the much larger hydrodynamic radius of the DNA.

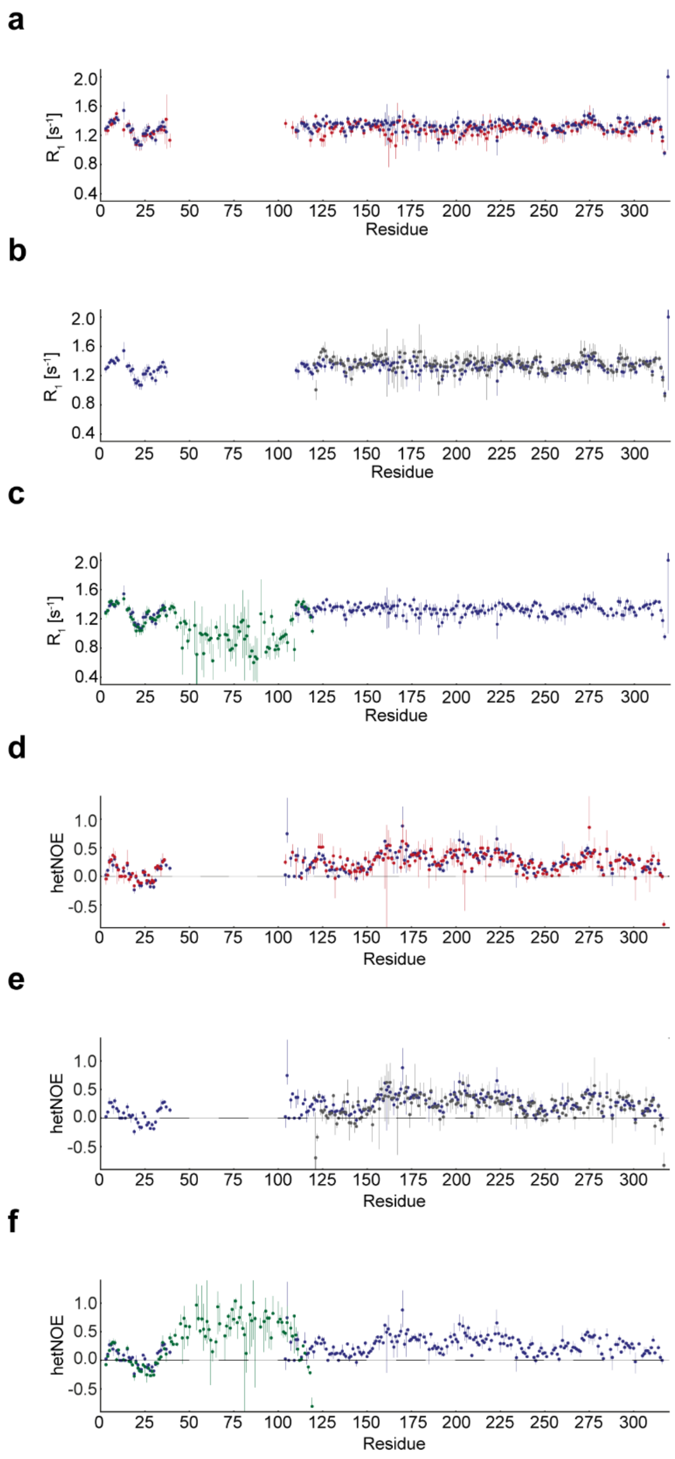


**Fig. S7. Relaxation data for full-length Sox2 and the isolated C-IDR, free and DNA-bound. a-c)** $R_1$ relaxation rates, and **d-f)** heteronuclear Overhauser effects (hetNOEs), for full-length Sox2 (blue), full-length Sox2 in complex with DNA (red), the isolated C-IDR (grey), and the isolated DBD (green).

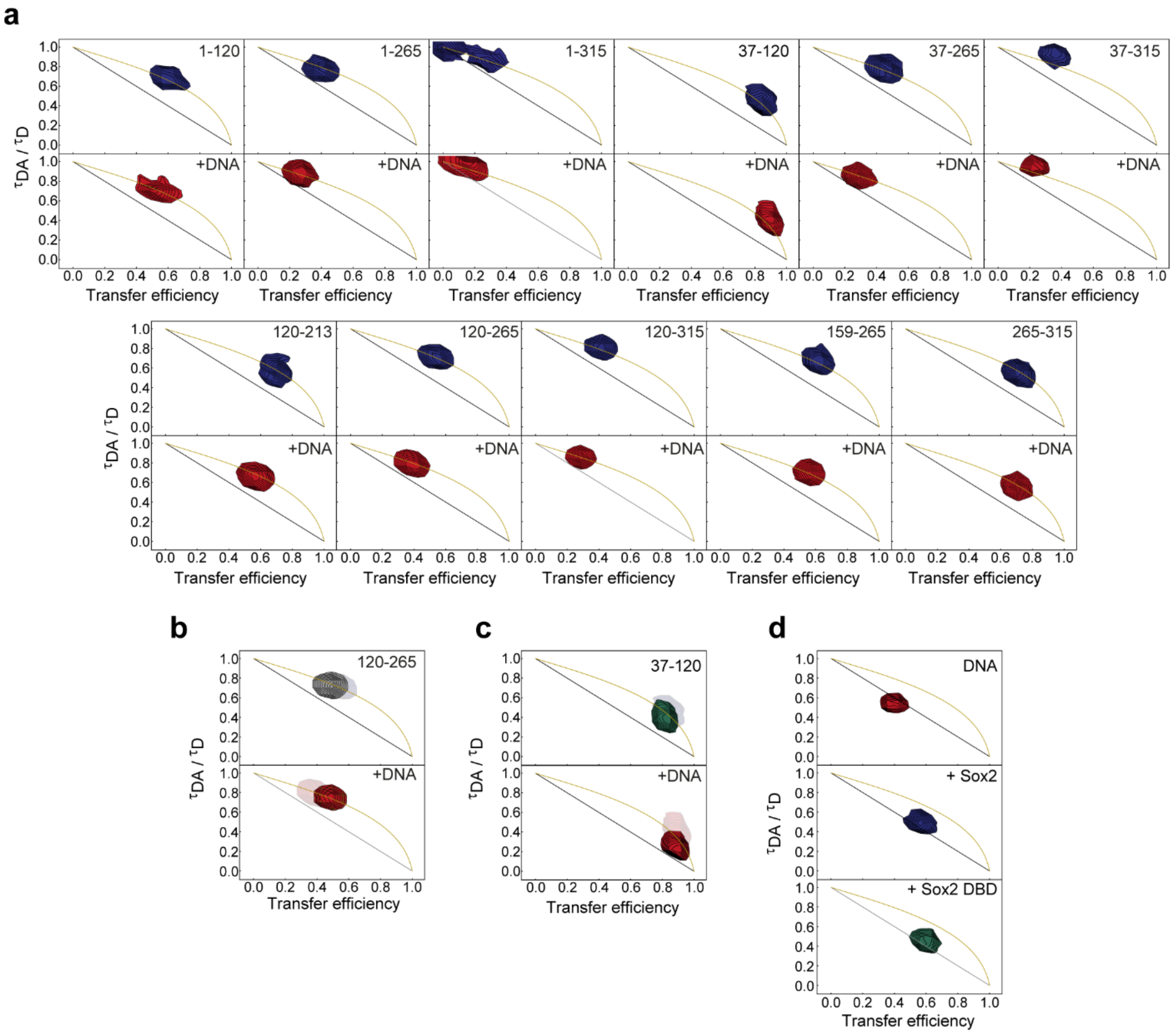


**Fig. S8. Fluorescence lifetime analysis of fluorescently labelled Sox2 variants and DNA. a)** Fluorescence lifetime analysis of all fluorescently labelled Sox2 variants, in the absence and presence of specific DNA. **b-c)** Fluorescence lifetime analysis of the isolated C-IDR (b) and DBD (c), in the absence and presence of DNA. The faint peaks represent the lifetimes of the full-length protein labelled in the same positions for comparison. **d)** Fluorescence lifetime analysis of fluorescently labelled 15 bp DNA in the free state (top), bound to full-length Sox2 (middle), and to the isolated DBD (bottom). Black lines describes the dependence of a static distance from the Förster equation, the yellow lines describe the dependence for a SAW-$\nu$ distance distribution[3].

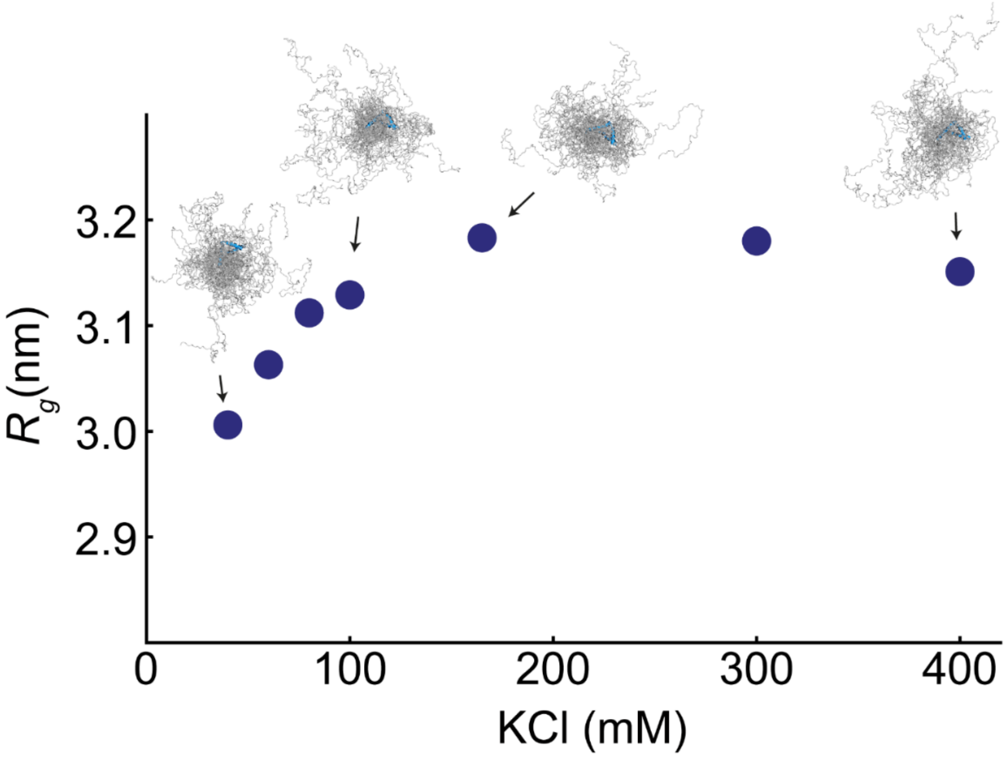


**Fig. S9. Radius of gyration calculated from simulations at different salt concentrations for free Sox2.** $R_g$ (for the region encompassing residues 120-265) as a function of apparent salt concentration for free Sox2 calculated from the simulations. At low salt the chain is compact (low $R_g$) and then expands due to increased charge screening with increasing salt concentrations, in agreement with FRET data (Fig. 2f). 20 snapshots from the simulations at different salt concentrations are shown.

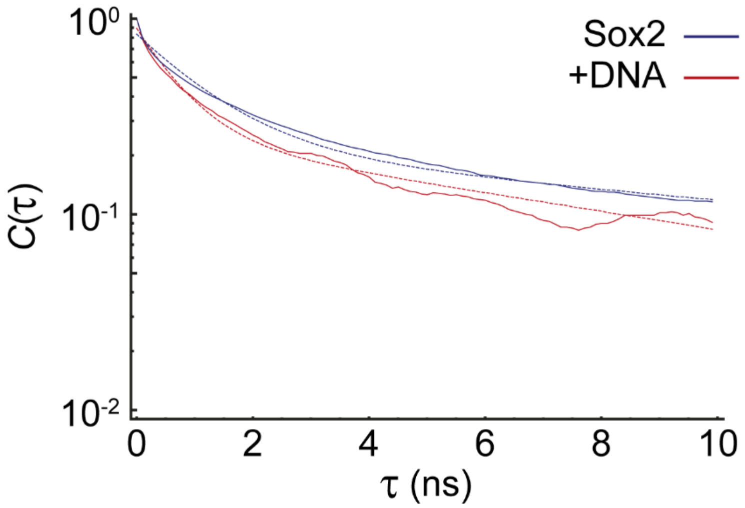


**Fig. S10. Contact lifetimes between the Sox2 Ser-rich domain and the DBD calculated from simulations.** The contact lifetimes, calculated from fitting the auto-correlation function using a double exponential function (dotted lines), are similar for free and DNA-bound Sox2 Ser-rich region.

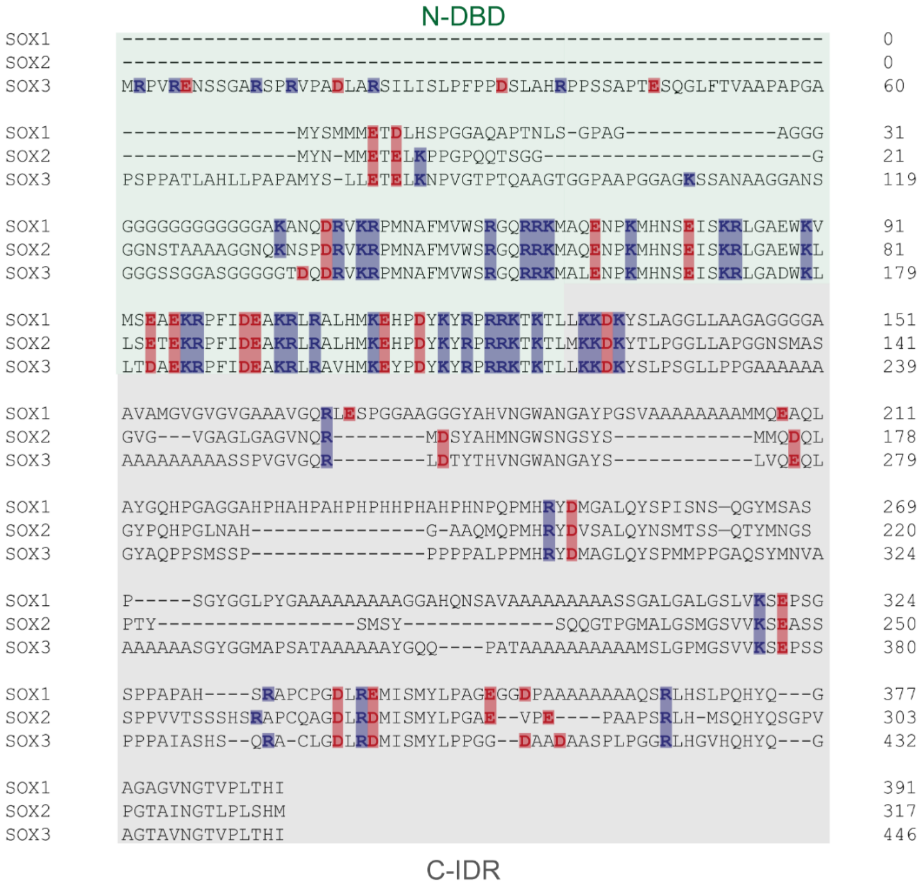


**Fig. S11. Sequence alignment of the SoxB family of TFs.** N-DBD and C-IDR regions are indicated with green and gray background, respectively. Charges are highlighted for clarity. Alignment generated by CLUSTAL O[4]. UniProt IDs: O00570 (Sox1), P48431 (Sox2) and P41225 (Sox3).

**Supplementary Table 1.** Sox2 variants used in this study. The residues used for fluorophore labelling (after substituting to cysteine) are marked in red.

| | |
|---|---|
| Full-length Sox2 | MYNMMETELKPPGPQQTSGGGGGNSTAAAAGGNQKNSPDRVKRPMNAFMVWSRGQRRKMAQENP KMHNSEISKRLGAEWKLLSETEKRPFIDEAKRLRALHMKEHPDYKYRPRRKTKTLMKKDKYTLPGGLLAPG GNSMASGVGVGAGLGAGVNQRMDSYAHMNGWSNGSYSMMQDQLGYPQHPGLNAHGAAQMQP MHRYDVSALQYNSMTSSQTYMNGSPTYSMSYSQQGTPGMALGSMGSVVKSEASSSPPVVTSSSHSRAP CQAGDLRDMISMYLPGAEVPEPAAPSRLHMSQHYQSGPVPGTAINGTLPLSHM |
| C-IDR | MKKDKYTLPGGLLAPGGNSMASGVGVGAGLGAGVNQRMDSYAHMNGWSNGSYSMMQDQLGYPQHP GLNAHGAAQMQPMHRYDVSALQYNSMTSSQTYMNGSPTYSMSYSQQGTPGMALGSMGSVVKSEASSS PPVVTSSSHSRAPCQAGDLRDMISMYLPGAEVPEPAAPSRLHMSQHYQSGPVPGTAINGTLPLSHM |
| N-DBD | MYNMMETELKPPGPQQTSGGGGGNSTAAAAGGNQKNSPDRVKRPMNAFMVWSRGQRRKMAQEN PKMHNSEISKRLGAEWKLLSETEKRPFIDEAKRLRALHMKEHPDYKYRPRRKTKTL |
| DBD | KNSPDRVKRPMNAFMVWSRGQRRKMAQENPKMHNSEISKRLGAEWKLLSETEKRPFIDEAKRLRALH MKEHPDYKYRPRRKTKTLMKK |

**Supplementary Table 2.** Measured FRET efficiencies and fluorescence lifetimes of all protein variants and DNA. The average donor and acceptor lifetimes are 2.72±0.11 ns and 3.01±0.04 ns, respectively.

| Sox2 variant | FRET efficiency (E) | Donor lifetime (ns) | Acceptor lifetime (ns) | FRET lifetime (ns) |
|---|---|---|---|---|
| 1-120 | 0.60 | 2.80 | 2.95 | 2.92 |
| 1-120 + DNA | 0.51 | 2.79 | 2.94 | 2.97 |
| 1-265 | 0.39 | 2.77 | 2.98 | 2.95 |
| 1-265 + DNA | 0.25 | 2.71 | 2.95 | 2.99 |
| 1-315 | 0.32 | 2.73 | 3.05 | 3.04 |
| 1-315 + DNA | 0.19 | 2.72 | 3.03 | 2.94 |
| 37-120 | 0.81 | 2.75 | 3.04 | 3.09 |
| 37-120 + DNA | 0.88 | 2.72 | 3.01 | 3.03 |
| 37-265 | 0.48 | 2.71 | 3.04 | 3.07 |
| 37-265 + DNA | 0.28 | 2.70 | 3.03 | 3.07 |
| 37-315 | 0.39 | 2.51 | 3.05 | 3.05 |
| 37-315 + DNA | 0.22 | 2.57 | 3.05 | 3.05 |
| 120-213 | 0.71 | 2.78 | 3.02 | 2.98 |
| 120-213 + DNA | 0.56 | 2.82 | 3.01 | 3.06 |
| 120-265 | 0.55 | 2.72 | 3.03 | 3.05 |
| 120-265 + DNA | 0.37 | 2.72 | 3.01 | 3.04 |
| 120-315 | 0.43 | 2.62 | 3.04 | 3.03 |
| 120-315 + DNA | 0.28 | 2.72 | 3.03 | 3.06 |
| 159-265 | 0.62 | 2.62 | 2.99 | 3.07 |
| 159-265 + DNA | 0.55 | 2.74 | 2.98 | 3.00 |
| 265-315 | 0.70 | 2.73 | 3.02 | 3.04 |
| 265-315 + DNA | 0.69 | 2.76 | 3.03 | 3.04 |
| C-IDR 120-265 | 0.48 | 2.62 | 3.01 | 3.08 |
| C-IDR 120-265 + DNA | 0.48 | 2.61 | 3.00 | 3.00 |
| DBD 37-120 | 0.81 | 2.46 | 3.00 | 2.98 |
| DBD 37-120 + DNA | 0.88 | 3.01 | 3.00 | 3.04 |
| Labelled 15 bp DNA | 0.40 | 2.89 | 3.11 | 3.11 |
| Labelled 15 bp DNA + Sox2 | 0.59 | 2.81 | 3.03 | 3.04 |
| Labelled 15 bp DNA + Sox2 DBD | 0.59 | 2.78 | 2.94 | 3.05 |

**Supplementary Table 3.** DNA constructs for binding experiments. The Sox2 binding site in the 601-Widom sequence is marked in red. 5AmMC6 indicates a C6-amino group for labelling with an NHS-ester fluorophore.

| | |
|---|---|
| 15 bp specific, (+) strand | 5'-/5AmMC6/ACT CTT TGT TTG GAT-3' |
| 15 bp specific, (-) strand | 3'-TGA GAA ACA AAC CTA/5AmMC6/-5' |
| 15 bp nonspecific, (+) strand | 5'-/5AmMC6/ TCT ATC TGT GTA TGT-3' |
| 15 bp nonspecific, (-) strand | 3'- AGA TAG ACA CAT ACA/5AmMC6/-5' |
| 30 bp specific, (+) strand | 5'-ATC CCA TTA GCA TCC AAA CAA AGA GTT TTC-3' |
| 30 bp specific, (-) strand | 3'-TAG GGT AAT CGT AGG TTT GTT TCT CAA AAG-5' |
| 21 bp specific, (+) strand | 5'-GAA TAC TCT TTG TTT GGA TGC-3' |
| 21 bp specific, (-) strand | 3'-CTT ATG AGA AAC AAA CCT ACG -5' |
| '601'-Widom sequence with Sox2 binding site at SHL +6, forward strand | 5'-TCCATGGACCCTATACGCGGCCGCCCTGGAGAATCCCGGTGCCGAGGCCGCTCAATTGGTCG TAGACAGCTCTAGCACCGCTTAAACGCACGTACGCGCTGTCCCCCGCGTTTTAACCGCCAAG GGGATTACTCCCTAGTCTCCAGGCCTTTGTTATGCAAATACATCCTGTGCATGTATTGAACA GCAGTATGCCT-3' |
| '601'-Widom sequence with Sox2 binding site at SHL +6, reverse strand | 3'-AGGCATACTGCTGTTCAATACATGCACAGGATGTATTTGCATAACAAAGGCCTGGAGACTAG GGAGTAATCCCCTTGGCGGTTAAAACGCGGGGGACAGCGCGTACGTGCGTTTAAGCGGTGC TAGAGCTGTCTACGACCAATTGAGCGGCCTCGGCACCGGGATTCTCCAGGGCGGCCGCGTAT AGGGTCCATGGA-5' |
| '601'-Widom primer, forward | 5'-/5AmMC6/TCCATGGACCCTATACGCGGCCGCC-3' |
| '601'-Widom primer, reverse | 3'-AGGCATACTGCTGTTCAATACATGCACAGGATGTATTTGCATAACAAAGGCCTGGAGAC/5AmMC6/-5' |

**Supplementary Table 4.** Binding affinities of Sox2 for DNA from smFRET experiments. Errors are standard errors based on propagating pipetting errors.

| **Binding reaction** | **$K_D$ (nM)** |
|---|---|
| Labelled specific DNA – Full-length Sox2 | 0.4 ± 0.2 |
| Labelled specific DNA – DBD | 0.3 ± 0.1 |
| Labelled Full-length Sox2 – 30 bp specific DNA | 0.7 ± 0.4 |
| Labelled DBD – 30 bp specific DNA | 0.3 ± 0.2 |
| Labelled non-specific DNA – Full-length Sox2 | 6 ± 2 |
| Labelled non-specific DNA – DBD | 11 ± 4 |

**Supplementary Table 5.** Contact lifetimes obtained from simulations for AD1, AD2, and Ser-rich domain, based on exponential fitting. Errors are the standard errors of the fit.

| | **$\tau$1 (ns)** | **$\tau$2 (ns)** |
|---|---|---|
| AD1 | 1.58 ± 0.03 | 18.12 ± 0.33 |
| AD1 + DNA | 1.32 ± 0.02 | 21.09 ± 0.27 |
| AD2 | 1.95 ± 0.03 | 24.34 ± 0.38 |
| AD2 + DNA | 0.48 ± 0.01 | 4.49 ± 0.06 |
| Ser-rich | 1.20 ± 0.02 | 16.09 ± 0.21 |
| Ser-rich + DNA | 0.71 ± 0.01 | 19.17 ± 0.122 |

**Supplementary video 1.** Simulation of free Sox2.

The free Sox2 ensemble, with the Sox2 DBD shown in blue and the disordered N-IDR and C-IDR in silver. The video shows 400 conformations of the Sox2 ensemble, collected from the 16 μs Langevin dynamics simulations (see Methods).

**Supplementary video 2.** Simulation of Sox2 in complex with DNA.

Ensemble of Sox2 bound to DNA, with the Sox2 DBD shown in blue, the disordered N-IDR and C-IDR in silver, and the DNA in dark grey. The video shows 400 conformations of the Sox2 ensemble in complex with DNA, collected from the 16 μs Langevin dynamics simulations (see Methods).
